# The RHIM within the M45 protein from murine cytomegalovirus forms heteromeric amyloid fibrils with RIPK1 and RIPK3

**DOI:** 10.1101/324590

**Authors:** Chi L. L. Pham, Merryn Strange, Ailis O’ Carroll, Nirukshan Shanmugam, Emma Sierecki, Yann Gambin, Megan Steain, Margaret Sunde

## Abstract

The M45 protein from murine cytomegalovirus protects infected murine cells from death by necroptosis and can protect human cells from necroptosis induced by TNFR activation, when heterologously expressed. We show that the N-terminal 90 residues of the M45 protein, which contain a RIP Homotypic Interaction Motif (RHIM), are sufficient to confer protection against TNFR-induced necroptosis. This N-terminal region of M45 drives rapid self-assembly into homo-oligomeric amyloid fibrils and interacts with the RHIMs of human RIPK1 and RIPK3 kinases to form heteromeric amyloid fibrils *in vitro*. An intact RHIM core tetrad is required for the inhibition of cell death by M45 and we show that mutation of those key tetrad residues abolishes homo- and hetero-amyloid assembly by M45 *in vitro*, suggesting that the amyloidogenic nature of the M45 RHIM underlies its biological activity. Our results indicate that M45 mimics the interactions made by RIPK1 with RIPK3 in forming heteromeric amyloid structures.

## Introduction

The RIP homotypic interaction motif (RHIM) was first identified in receptor-interacting protein kinase 1 (RIPK1) and receptor-interacting kinase 3 (RIPK3)^1^. A RHIM contains 18– 19 amino acids, with a tetrad (V/I)-Q-(V/I/L/C)-G core sequence (Fig. 1a). An intact RHIM is required for the interaction between RIPK1 and RIPK3 that occurs downstream of tumour necrosis factor receptor 1 (TNFR1) activation, during the programmed cell death response known as necroptosis^2,3^. In addition to the TNFR1 pathway that leads to RIPK1:RIPK3 association, two other pathways involving RIPK3 and specific RHIM-containing adapter proteins can result in necroptosis^4^. Binding of microbial pathogen-associated molecular patterns to Toll-like receptors 3 and 4 can lead to activation of RIPK3 through formation of a complex with the RHIM-containing TIR-domain-containing adapter-inducing interferon-β protein (TRIF). Additionally, the Z-DNA binding protein/DNA-dependent activator of interferon regulatory factors (ZBP-1/DAI) is a cytosolic nucleic acid sensor that binds foreign nucleic acid and acts as the RHIM adapter protein for RIPK3 in viral-induced necroptosis^5-8^. In all three pathways, oligomerisation and autophosphorylation of RIPK3 occurs after binding of RIPK3 to the adapter protein. The activation of RIPK3 leads to phosphorylation and oligomerisation of mixed lineage kinase domain-like protein (MLKL) and this commits the cell to membrane disruption, ion dyshomeostasis and lytic necrotic cell death^9-12^.

**Figure 1:**
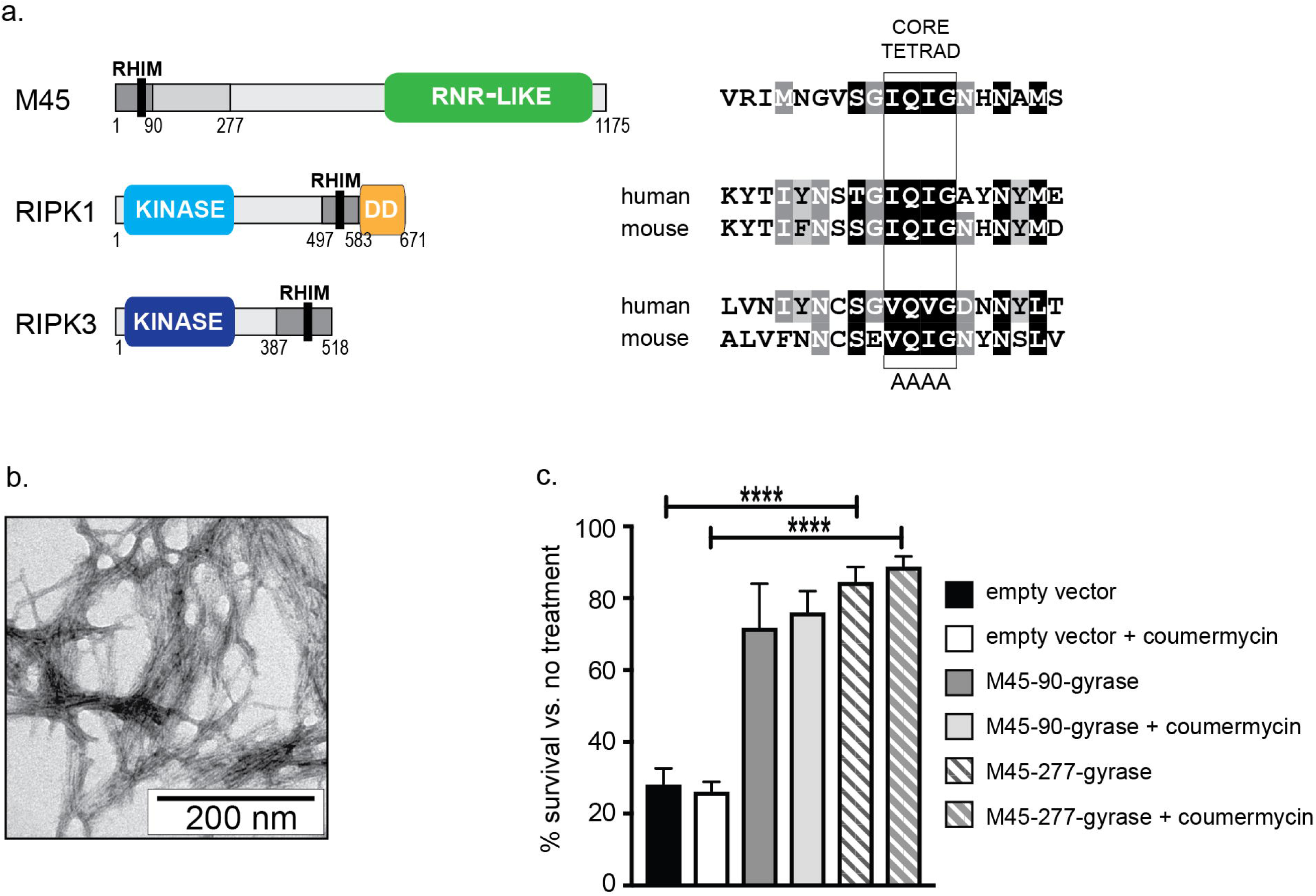
The M45 RHIM is amyloidogenic and 1-90 and 1-277 regions of M45 confer protection against TNF-induced necroptosis in human cells. (**a**) Schematic representation of M45, RIPK1 and RIPK3 proteins. Grey regions indicate RHIM-encompassing fragments used in this study. Black bar represents 19-residue RHIM. RNR-like represents inactive ribonucleotide reductase domain, DD represents the RIPK1 death domain. Amino acid sequences of the RHIMs of M45 and human and mouse RIPK1 and RIPK3, boxed region indicates the core tetrad that is substituted with AAAA in mutant protein constructs. (**b**) Negatively-stained transmission electron micrograph of fibrils formed by a synthetic peptide encompassing the M45 RHIM. Three samples were examined by microscopy. (**c**) HT29 cell survival 18h after treatment with TNF+BV-6+z-VAD-fmk to induce necroptosis. Cells expressed lentivirus constructs encoding 1-90 or 1-277 fragments of M45 and were cultured +/-the addition of coumermycin A1 to induce dimerisation. N=2 for M45-90 gyrase and 3 for M45-277 gyrase. Error bars represent SEM, **** = p<0.0001.

Necroptosis has essential functions in adult tissue homeostasis and innate immune defence against intracellular pathogens, however undesirable necroptosis can be triggered by ischaemic tissue damage^13,14^ and recent evidence suggests that necroptosis contributes to neurodegenerative conditions^15^. Small molecules that block necroptosis are under investigation as potential modulators of ischaemia-reperfusion injury associated with myocardial infarction, stroke and solid organ transplantation^16^. Solid cancers present with necrosis but additionally necroptosis can promote tumour cell growth through suppression of the immune response against cancer^17^. A number of herpesviruses express RHIM-containing proteins that are primarily implicated in inhibiting host cell necroptosis to ensure virus survival^18,19^. Understanding the molecular basis for this viral RHIM-driven inhibition could identify therapeutic targets and provide guidance for the development of anti-necroptosis strategies with many clinical applications.

We have characterised the properties of the RHIM within the protein M45 (Fig. 1a), also known as the viral Inhibitor of RIP Activation (vIRA)^20^ from murine cytomegalovirus (MCMV) and the structural basis for its interactions with RIPK1 and RIPK3. MCMV inhibits host apoptosis and necroptosis pathways to sustain infection^21,22^ and specifically thwarts necroptosis in infected endothelial cells and macrophages^23,24^ through the actions of M45. Thus far, M45 is the only viral RHIM-containing protein known to suppress necroptosis in mouse and human cells^23,25,26^. The M45 protein is delivered within the infecting virions^27^. Inhibition of necroptosis by M45 has been shown to involve interactions between M45 and RIPK1^23^, RIPK3^26^ and ZBP-1/DAI^7,28^ and to require an intact RHIM; substitution of the IQIG tetrad within the RHIM of M45 with AAAA abolishes the ability of the virus to confer protection against necroptosis^7,28^.

In 2012 Wu and colleagues demonstrated that the RHIM-driven association between RIPK1 and RIPK3 results in the formation of an amyloid, fibrillar structure where the RHIM residues form the β-sheet core of RIPK1:RIPK3 fibrils^29^. This necrosome complex was the first functional amyloid structure described with an active signalling role^30^. The report of a similar amyloid-based signalling platform in *Drosophila*, where the PGRP-LC and PGRP-LE receptors and downstream Imd adapter protein interact through motifs bearing some similarity to RHIMs^31^, suggests that RHIM-based signalling may be widespread. The structure of the necrosome RIPK1:RIPK3 core has been determined by solid-state nuclear magnetic resonance spectroscopy (ssNMR), confirming the hetero-amyloid nature of the complex and revealing a compact hydrophobic interface involving the RHIM tetrad^32^.

Given the demonstrated ability of the M45 protein to protect against TNFR-initiated necroptosis in human cells^25^ and the interest in inhibition of necroptosis in clinical settings, we have investigated the interactions between the RHIMs of M45 and human RIPK1 and RIPK3, which are highly homologous to the murine RIPKs (Fig. 1a). We find that the RHIM of M45 is itself amyloidogenic and the ability of M45 to self-assemble into fibrils depends on the presence of the core tetrad sequence IQIG. A 90-residue fragment of M45 encompassing the RHIM is sufficient to confer protection against TNF-driven necroptosis in human cells. The RHIM of M45 is readily incorporated into heteromeric amyloid fibrils with the minimum portions of human RIPK1 or RIPK3 that are necessary for necrosome formation^29^. The viral M45 RHIM preferentially assembles with RIPK3 over RIPK1 and based on this, we have generated a model of a hetero-amyloid structure with RIPK3. We observe that incorporation of the viral M45 RHIM alters the properties of the human RHIM-based complexes. These results suggest that the M45 protein sequesters host RHIM-containing proteins in alternative amyloid structures that are incompatible with auto-phosphorylation of RIPK3 and/or activation of the downstream necroptosis mediator MLKL. The ability of the M45 protein revealed here, to form amyloid-based structures with host RHIM-containing proteins, suggests a general mechanism for modulation of host RHIM-based signalling by other viral RHIM-containing proteins.

## Results

### The M45 RHIM drives spontaneous amyloid fibril formation

Given the discovery of the amyloid nature of the RIPK1;RIPK3 necrosome^29^, we initially sought to determine whether the RHIM in M45 is amyloidogenic. A peptide of 19 residues, corresponding to the M45 RHIM sequence, readily assembled into long fibrils with amyloid morphology (Fig. 1a, b). In order to investigate the nature of the interactions between M45 and host proteins, we prepared fusion proteins containing portions of M45, RIPK1 and RIPK3 (Fig. 1a) with different partner proteins (His_6_-ubiquitin (His-Ub), His_6_-YPet, -ECFP or -mCHERRY fluorescent proteins (FPs) or maltose binding protein (MBP)). The human RIPK protein constructs contained the minimum RHIM-containing regions shown to support interactions between these two proteins, i.e. RIPK1-(497-583) and RIPK3-(387-518)^29^. Two RHIM-encompassing fragments of M45 were tested, comprising 90 or 277 N-terminal residues of the viral protein. The 1–90 region of M45 was identified as the likely smallest autonomous fragment of M45 because analysis of the predicted secondary structure of the protein highlighted a clear boundary between the β-sheet elements flanking the RHIM and the strongly predicted helical elements from residue 93 onwards. The 277 N-terminal residues of M45 have previously been shown to provide protection against some forms of cell death^23^ and to inhibit ZBP-1 signalling^7^.

### RHIM-encompassing fragments of M45 inhibit necroptosis in human HT-29 cells

We compared the ability of the 1–90 and 1–277 regions of M45 to protect against TNFR-induced necroptosis in human HT-29 cells. The 90-residue and 277-residue portions were expressed as fusion proteins with the B subunit of *E. coli* DNA gyrase^9^. Addition of the antibiotic coumermycin A1 can induce dimerization of the gyrase subunit, to substitute for the potential dimerisation of the inactive ribonucleotide reductase-like (RNR1-like) domain of M45, which may form a heterodimer with RNR large and small subunits in the intact viral protein. Fig. 1c shows that the 90-residue and 277-residue portions of M45 protect human cells from TNFR-induced necroptosis through the RIPK1:RIPK3 pathway to a similar extent, in the range 70–90%. This is independent of the addition of coumermycin A1, indicating that dimerisation of M45 is not necessary for its role in inhibition of host cell death. For the majority of biophysical experiments we have therefore probed RHIM-driven interactions using the minimum necrosome-forming regions of RIPK1 and RIPK3^29^ and the 90-residue N-terminal portion of M45, which displays the ability to inhibit TNFR-induced necroptosis.

### The wild type core tetrad is critical for amyloid formation by M45 and RIPK1 but not RIPK3

The RHIM-containing regions were expressed as fusion proteins with His-Ub or His-FP to provide a means to study the kinetics of hetero-amyloid assembly. These fusion proteins were purified and maintained in the presence of denaturant, to prevent unwanted self-assembly. Prior experience with other functional amyloid-forming proteins has demonstrated that upon dilution of the denaturant, the ubiquitin domain refolds independently to its native state, allowing the characteristics of the partner sequence to be studied^33^. In YPet, ECFP and mCHERRY-containing RHIM fusion proteins, the fluorescent partner refolded correctly, as judged by recovery of characteristic fluorescence profiles. Purified His-Ub-M45_1-90_ spontaneously formed fibrils upon removal of denaturant by dialysis and the fibrils displayed a cross-β X-ray fibre diffraction pattern, with a strong, sharp meridional reflection at 4.7 Å and a weaker, diffuse equatorial reflection at ∼10.5 Å, which are characteristic of amyloid fibrils^34^ (Fig. 2a).

**Figure 2:**
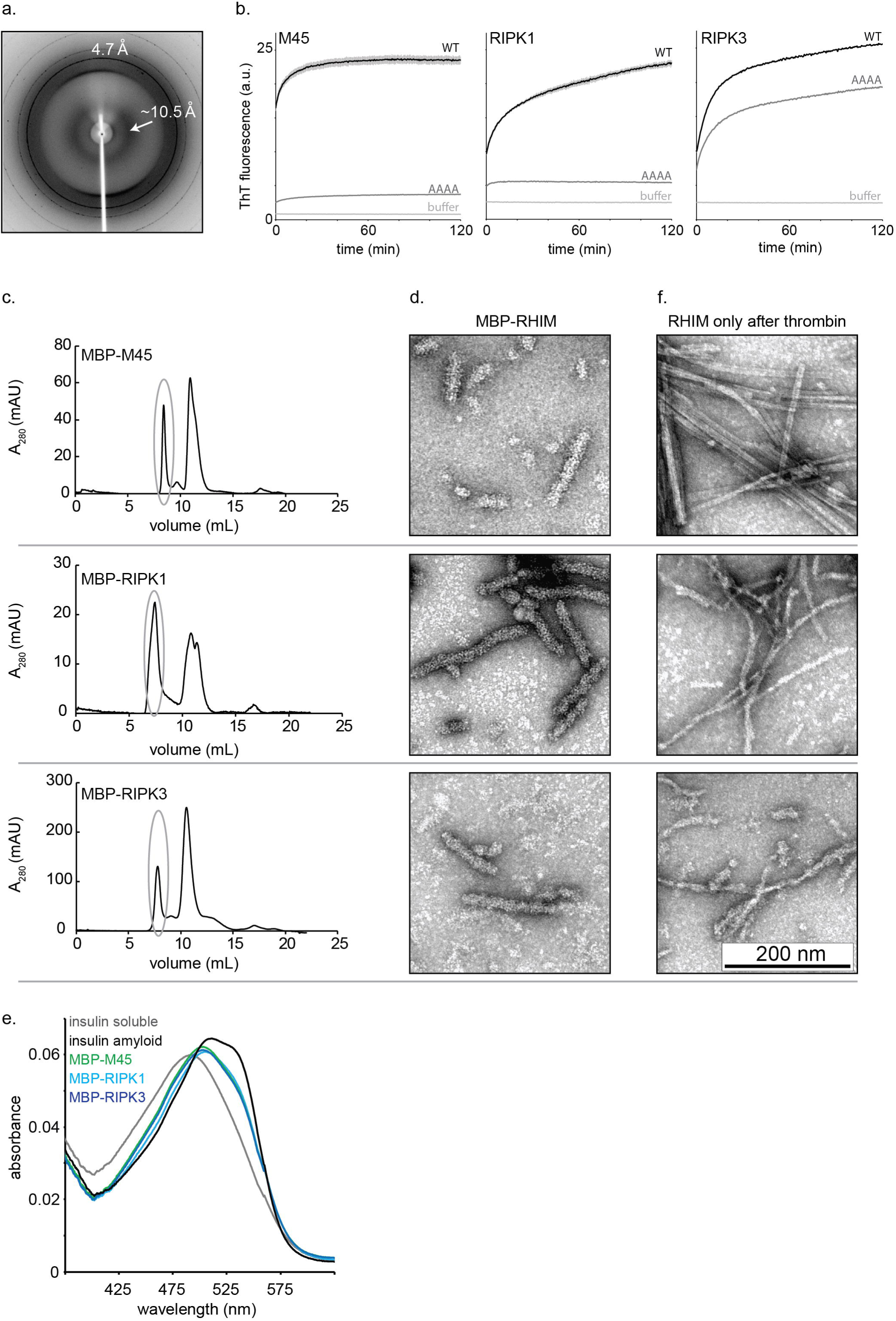
The RHIM-containing fragment forms the amyloid fibril core. (**a**) X-ray fibre diffraction pattern collected from fibrils formed by His-Ub-M45_1-90_. Two different samples were examined by XRFD. (**b**) Thioflavin T fluorescence as a function of time, following dilution of WT or AAAA His-Ub-M45_1-90_, mCHERRY-RIPK1_497-583_, and YPet-RIPK3_387-518_ from 8 M-containing buffer to 100–300 mM urea. Buffer sample contains ThT but no protein. Samples were tested in triplicate and solid line is the average of the triplicates and the error bars (in grey) represent +/-SEM. (**c**) Size exclusion chromatography (SEC) of MBP-M45_1-90_, MBP-RIPK1_497-583_, and MBP-RIPK3_387-518_ following amylose affinity purification. Grey oval indicates high MW material eluting in void volume. (**d**) Negatively-stained transmission electron micrograph of material eluted in the void volume. These are representative micrographs of the material observed. All fusion proteins expressed and purified at least twice. (**e**) Absorbance spectrum from solutions containing Congo red with soluble monomeric insulin, insulin amyloid fibrils or material eluted in the void volume containing MBP-M45_1-90_ (green), MBP-RIPK1_497-583_ (cyan), or MBP-RIPK3_387-518_ (blue). Experiment conducted once. (**f**) Representative images of insoluble material generated after treatment with thrombin removed the MBP. Experiment repeated three times.

The kinetics of self-assembly, or homo-oligomeric fibril formation, of the RHIM-containing fusion proteins were studied by monitoring the fluorescence of thioflavin T (ThT), a dye commonly used as an indicator of amyloid structure. ThT exhibits increased fluorescence emission at 485 nm when it binds to amyloid fibrils^35^. All wild type RHIM-containing His-Ub and fluorescent fusion proteins formed homomeric amyloid (Fig. 2b). Tetra-alanine substitution of the RHIM core is a commonly used strategy to probe the role of RHIM:RHIM interactions in cells and during viral infection^1,23^ (Fig. 1a). RHIM mutant constructs in which the four core amino acids were substituted by AAAA were produced and the ability of these proteins to assemble into homomeric amyloid fibrils was tested. The substitution of the IQIG tetrad by AAAA within the M45_1-90_ and RIPK1_497-583_ RHIM constructs abolishes amyloid assembly (Fig. 2b). In contrast, the AAAA version of RIPK3_387-518_ retains the ability to assemble into amyloid fibrils and has a similar kinetic profile to the WT form but exhibits a reduced ThT fluorescence intensity, reflecting an alteration in the fibril structure. This result indicates that in RIPK3, interactions outside of the core four residues are also important in the formation of homomeric amyloid structures.

### The M45 RHIM forms the core structure within the amyloid fibrils

Amyloid assembly by the RHIM-containing fragments was also characterised under native conditions, when amyloid assembly occurred spontaneously during recombinant expression. When M45_1-90_, RIPK1_497-583_ and RIPK3_387-518_ were expressed as fusions with MBP, the fusion proteins remained soluble and could be purified under native conditions. Size exclusion chromatography after affinity purification separated the material into two main fractions (Fig. 2c). The first of these eluted in the void volume, indicating the formation of structures with MW >100 000 Da. SDS PAGE showed that the peak eluting in the void volume contained MBP-RHIM fusion protein while the second peak contained MBP only. The high molecular weight material was examined by negative stain transmission electron microscopy (TEM) and for MBP-M45_1-90_, MBP-RIPK1_497-583_ and MBP-RIPK3_387-518_, fibrillar structures of up to ∼200 nm in length were observed (Fig. 2d). Congo red solution binding assays performed with this material showed increased absorbance at 540 nm for all three samples and a shift in the absorbance maxima, relative to a non-amyloid control (soluble insulin), indicating that the fibrils had an amyloidogenic nature (Fig. 2e).

To probe the arrangement of the RHIM within the fibrils, we treated the fibrillar material with thrombin to cleave specifically between MBP and the RHIM-containing regions. During incubation with the protease, insoluble material became visible in all three samples. Separation of this material from the soluble fraction by centrifugation, followed by SDS PAGE and SEC analysis of the soluble material, revealed that M45_1-90_, RIPK1_497-583_ and RIPK3_387-518_ had formed insoluble precipitates while the MBP remained soluble. The insoluble material was resuspended and examined by TEM and fibrils were observed that were longer and thinner than those formed by the MBP fusion proteins (Fig. 2f). These results indicate that the RHIM-containing region of M45 is amyloidogenic and forms the β-sheet core structure of the fibrils, in a similar fashion to that demonstrated for RIPK1 and RIPK3 by Wu and coworkers^29^. The MBP fusion partner is arrayed on the outside of the β-sheet core and limits the growth of the fibrils to ∼200 nm; in its absence, the RHIM-containing segments assemble into longer fibrils.

### M45 interacts with RIPK1 and RIPK3 to form heteromeric assemblies

To address the possibility of hetero-amyloid formation involving the RHIM-containing portions of M45, RIPK1 and RIPK3, we applied fluorescence polarisation and co-localisation experiments. When His-Ub-RIPK1_497-583_ was covalently labelled with Alexa Fluor488 and mixed with unlabelled His-Ub-RIPK1_497-583_, His-Ub-RIPK3_387-518_ or His-Ub-M45_1-90_ in a ratio of 1 labeled:1999 unlabelled, the Alexa488-labelled RIPK1_497-583_ was incorporated into growing His-Ub-RIPK1_497-583_, His-Ub-RIPK3_387-518_ or His-Ub-M45_1-90_ fibrils (Fig. 3a). The rates of incorporation of the RIPK1-RHIM construct into His-Ub-RIPK1_497-583_ and His-Ub-RIPK3_387-518_ fibrils were similar and RIPK1-RHIM was incorporated into His-Ub-M45_1-90_ fibrils more slowly. The tetrad sequence in the RHIM of M45 is IQIG and identical to that in RIPK1, whereas the RIPK3 tetrad sequence is VQVG. The observed difference in incorporation rates indicates that residues outside the tetrad also influence the interactions between RHIM-containing proteins (Fig. 1a). Taken together with the observed different effects of the AAAA mutations on M45, RIPK1 and RIPK3 homomeric amyloid formation, we conclude that the contributions of the tetrad sequences to homomeric and heteromeric structures are distinct and differ between RHIM-containing proteins.

**Figure 3:**
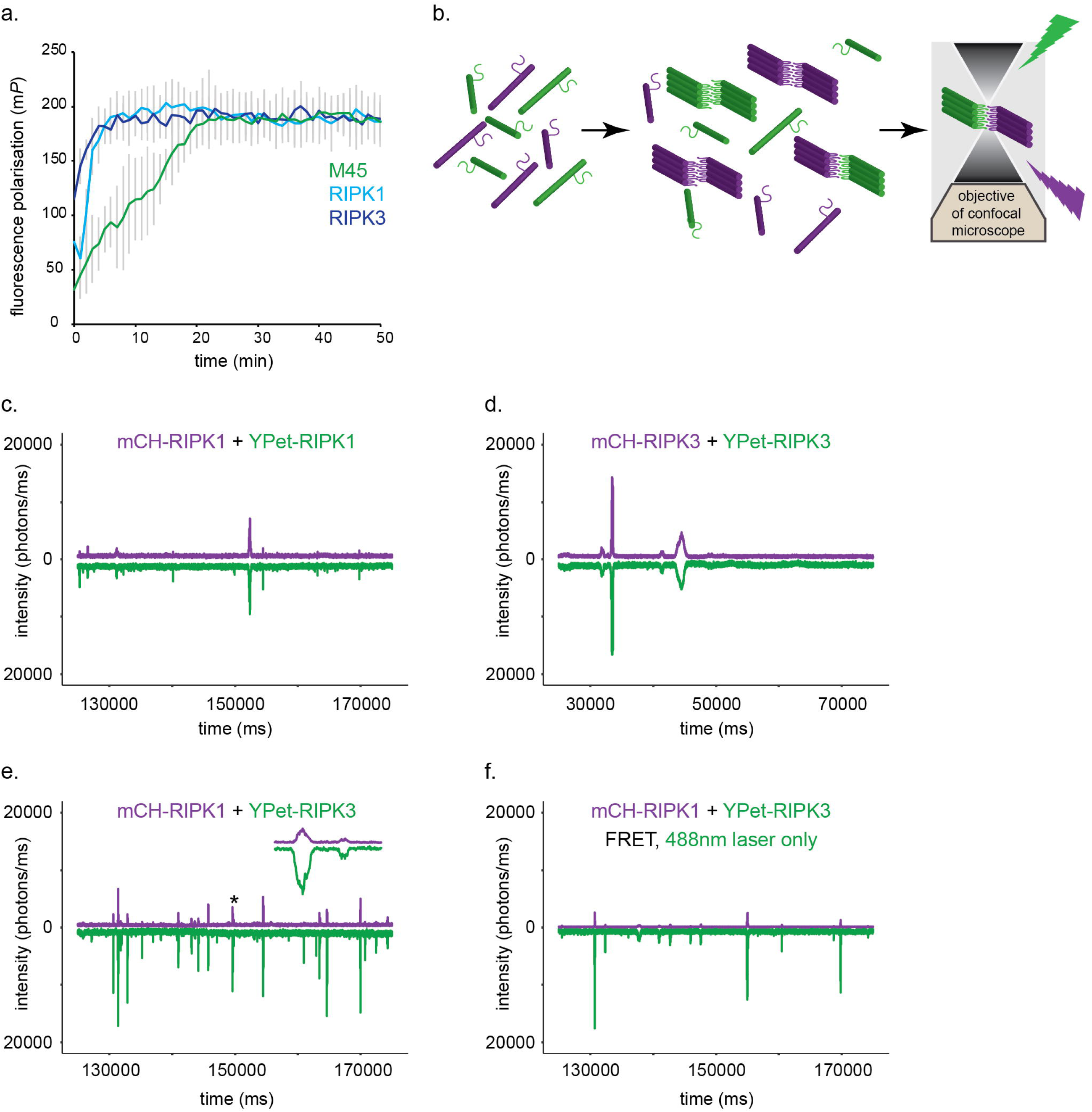
Assembly of RHIMs gives rise to homomeric and heteromeric structures. **(a)** Fluorescence polarization profiles indicating the rate of incorporation of Alexa488-tagged RIPK1_497-583_ into unlabeled M45_1-90_ (green), RIPK1_497-583_ (cyan), or RIPK3_387-518_ (blue), with labeled:unlabeled protein at 1:1999. The average polarization profile from triplicate readings is shown, with s.d. indicated. (**b**) Schematic representation of fluorescence spectroscopy experiments used to detect the formation of homo- and heteromeric amyloid fibrils. Fusion proteins containing RHIMs and different fluorescent partners are mixed together under assembly-permissive conditions and examined in the confocal volume. Co-assembly of two different fluorophores within one complex results in detection of coincident signals in the two channels. (**c**) Representative fluorescence intensity trace showing homomeric assembly over 40 s in a mixture of mCHERRY-RIPK1_497-583_ and YPet-RIPK1_497-583_. (**d**) Representative fluorescence intensity trace showing homomeric assembly over 40 s in a mixture of mCHERRY-RIPK3_387-518_ and YPet-RIPK3_387-518_. (**e**) Representative fluorescence intensity trace showing heteromeric assembly over 40 s in a mixture of mCHERRY-RIPK1_497-583_ and YPet-RIPK3_387-518._ Inset is expansion of region indicated by * on main trace. (**f**) Förster resonance energy transfer (FRET) signal between YPet- and mCHERRY fluorophores, detected with excitation by 488 nm laser only. All mixtures examined in at least three different experiments on three separate occasions. In panels c–f the signal from the mCHERRY-tagged protein is coloured magenta and the signal from the YPet-tagged protein is coloured green.

Fluorescence co-localisation experiments were applied to detect RHIM-based hetero-oligomers against a background of monomeric proteins and the potential competing formation of homo-oligomers^36,37^. In these experiments, multiple different RHIM-containing fusion proteins were mixed together and maintained in monomeric form in 8 M urea-containing buffer. Assembly or co-assembly were initiated by reduction of the urea concentration through dilution. Mixtures of YPet- and mCHERRY-tagged proteins were examined in solution, using a confocal microscope where the lasers create a very small confocal volume (∼250×250×800 nm). Freely diffusing fluorophore-tagged proteins are only excited and detected when they are within the confocal volume (Fig. 3b). The number of molecules in the focal volume/millisecond over the range of concentrations used was ∼10-500.

In initial fluorescence spectroscopy experiments, we analysed mixtures of YPet- and mCHERRY-tagged RIPK1_497-583_ or YPET- and mCHERRY-tagged RIPK3_387-518_. We observed coincidence of the signals from both fluorophores when oligomers formed, demonstrating that homomeric fibril assembly can be detected in this way (Fig. 3c,d). As expected, mixtures of mCHERRY-RIPK1_497-583_ and YPet-RIPK3_387-518_ exhibited formation of heteromeric fibrils, indicated by coincidence of the signals from the two fluorophores attached to different RHIM-containing constructs (Fig. 3e). In addition, FRET could be observed between YPet and mCHERRY within the fibrils (Fig. 3f). These results provide the first direct single-molecule-level evidence for the formation of heteromeric fibrils containing RIPK1 and RIPK3 RHIM sequences, supporting the results from bulk solution previously reported^29^. The RIPK3_387-518_ construct demonstrated a greater propensity to self-assemble than the RIPK1_497-583_ construct, as evidenced by the formation of larger assemblies (Fig. 3c,d). In addition, all oligomeric RIPK1_497-583_ appeared coincident with RIPK3_387-518_ material, while some RIPK3_387-518_-only fibrils were observed (Fig. 3e).

The self-assembly of M45_1-90WT_ into fibrils could be detected by fluorescence co-localisation and, as expected given the ThT results, substitution of the IQIG tetrad with AAAA abolished the ability of this M45 fragment to self-assemble (Fig. 4a,b). This was confirmed by electron microscopy, which detected the presence of long fibrils formed by M45_1-90WT_-mCHERRYand only amorphous clusters in the M45_1-90AAAA_-mCHERRY sample (Supplementary Fig. 1).

**Figure 4:**
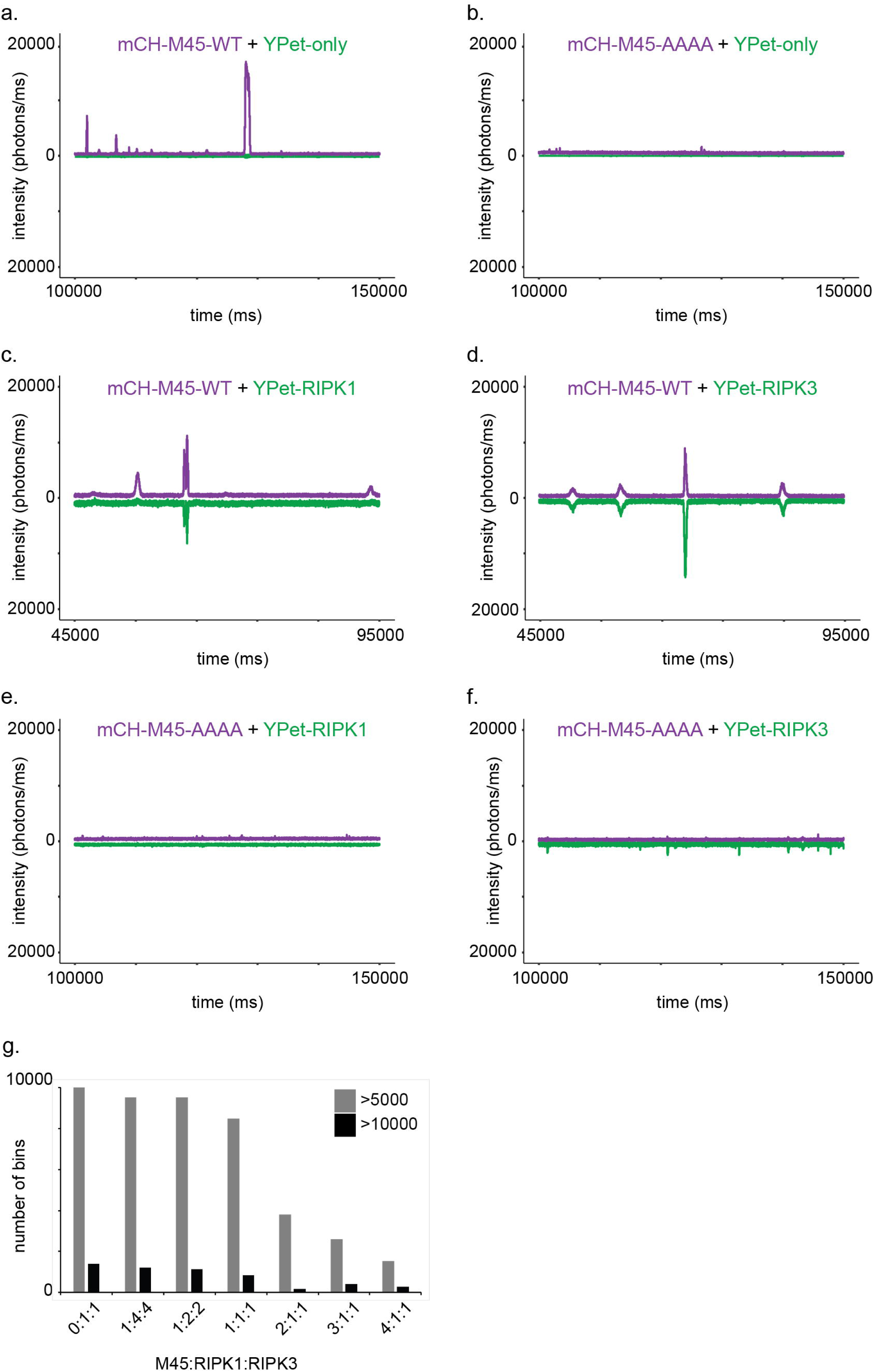
Heteromeric amyloid formation including M45 RHIM requires intact core tetrad. Representative fluorescence intensity traces collected over 50 s for mixtures of (**a**) mCHERRY-M45_1-90WT_ and YPet-only; (**b**) mCHERRY-M45_1-90AAAA_ and YPet-only; (**c**) mCHERRY-M45_1-90WT_ and YPet-RIPK1_497-583_; (**d**) mCHERRY-M45_1-90WT_ and YPet-RIPK3_387-518_; (**e**) mCHERRY-M45_1-_ 90AAAA and YPet-RIPK1_497-583_ and (**f**) mCHERRY-M45_1-90AAAA_ and YPet-RIPK3_387-518._ (**g**) Number of bins where intensity (photons/ms) exceeded 5000 or 10000 in the last 200 s of a 10–min incubation, where samples contained no His-Ub-M45_1-90_ or up to a 4-fold molar excess of His-Ub-M45_1-90_. Coincidence experiments conducted three times or more on different days. Quantification performed once.

When M45_1-90WT_-mCHERRY and YPet-RIPK1_497-583_ or YPet-RIPK3_387-518_ were mixed under fibril assembly conditions, we observed coincidence of M45_1-90WT_ with both RIPK fragments, demonstrating heteromeric complex formation between viral and host RHIM-containing fragments (Fig. 4c,d). The complexes containing M45_1-90_ and RIPK3_387-518_ appeared larger than those containing M45_1-90_ and RIPK1_387-518_, as judged by the length of time taken by the heteromeric complexes to diffuse through the focal volume. The AAAA mutation of the core tetrad of M45 abolished co-assembly with RIPK1 and RIPK3 (Fig. 4e,f). This result suggests that the ability of the M45 RHIM to inhibit host necroptosis is associated with its ability to form heteromeric amyloid structures with RIPK1 and RIPK3.

We added non-fluorescent His-Ub-M45_1-90_ into mixtures of YPet- and mCHERRY-tagged RIPK1 and RIPK3 fragments, to probe the effect of the amyloidogenic viral RHIM on heteromeric complex formation by the two human RHIMs. The number of times a large complex generating >5000 or >10000 photons above baseline intensity was detected in the focal volume for 1 millisecond during the last 200 seconds of a 10–minute incubation was quantified for a RIPK1:RIPK3 mixture and a series of RIPK1:RIPK3:M45 mixtures. The addition of His-Ub-M45_1-90_ resulted in a decrease in the number of the large complexes detected, correlated with the relative concentration of His-Ub-M45_1-90_ (Fig. 4g). EM of samples formed with increasing amounts of M45_1-90_ relative to RIPK3_387-518_ (1:1; 2:1 and 4:1) confirmed an increase in the size of the fibrillar assemblies (Supplementary Fig. 2). The reduction in the number of large complexes detected is therefore likely due to a reduction in the rate at which larger complexes diffuse in the well, which results in fewer passes through the focal volume and hence detection. Collectively, these results show that M45_1-90_ is able to interact with RIPK1_497-583_ and RIPK3_387-518_ in a RHIM-dependent manner but displays a preference for co-assembly with the RHIM of RIPK3.

### The M45-RHIM alters the formation of fibrils by the RHIMs of RIPK1 and RIPK3

Self-assembly of single types of RHIM-containing proteins resulted in predominantly unbranched single fibrils, as visualised by TEM (Fig. 5a). In contrast, when the M45_1-90_ fragment was mixed with either of the RIPK1_497-583_ or RIPK3_387-518_ fragments in a 1:1 molar ratio under conditions allowing fibril assembly, extensive fibrillar networks were observed (Fig. 5a). These consisted of large bundles of multiple fibrils that branched and reformed in different combinations, resulting in an interweaved and dense assembly covering large areas of the TEM grids. By contrast, when RIPK1_497-583_ and His-Ub-RIPK3_387-518_ were mixed together under conditions allowing fibril assembly, isolated mostly single fibrils of <200 nm length were observed, similar to those reported by Li et al^29^. When M45_1-90_ or RIPK1_497-583_ were allowed to assemble in the presence of an unrelated protein with the same fusion partner, His_6_-Ub-RodA, no thick multi-fibril bundles were observed, suggesting that this is a feature associated with hetero-fibrils formed by two RHIM-containing proteins (Supplementary Fig. 3).

**Figure 5:**
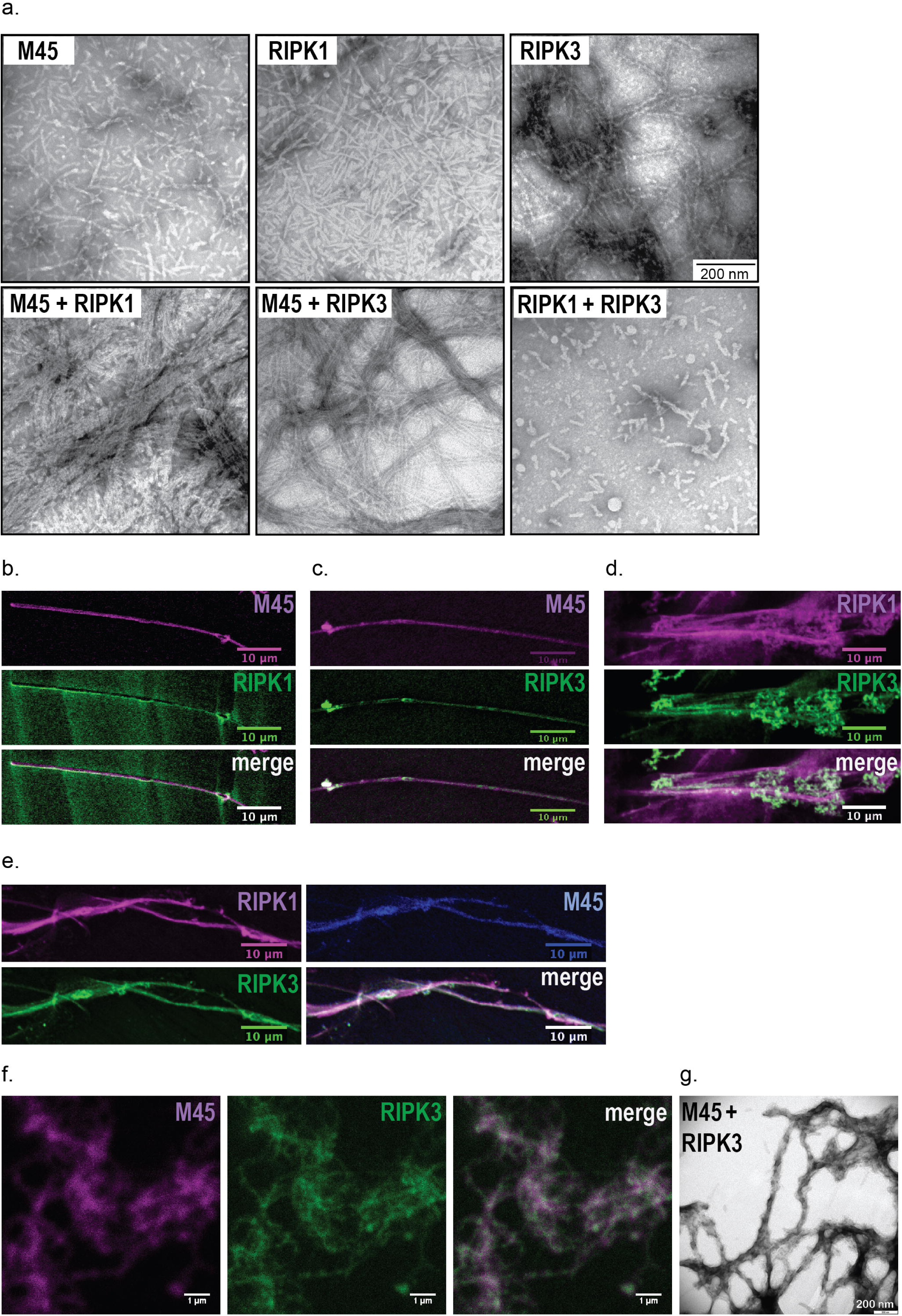
Co-assembly of M45_1-90WT_ with RIPK1_497-583_ and His-Ub-RIPK3_387-518_ results in the formation of heteromeric fibrils and dense fibrillar networks. (**a**) Representative electron micrographs of fibrils formed by M45_1-90_, RIPK1_497-583_ or His-Ub-RIPK3_387-518_ when dialysed individually or in mixtures, into phosphate buffer to remove urea. Confocal fluorescence images of heteromeric fibrils formed by co-assembly of (**b**) mCHERRY-M45_1-90_ and YPet-RIPK1_497-583_; (**c**) mCHERRY-M45_1-90_ and YPet-RIPK3_387-518_; (**d**) mCHERRY-RIPK1_497-583_ and YPet-RIPK3_387-518_; (**e**) ECFP-M45_1-90_ with mCHERRY-RIPK1_497-583_ and YPet-RIPK3_387-518_. (**f**) STED super-resolution and (**g**) electron microscope images of heteromeric fibrils formed by co-assembly of mCHERRY-M45_1-90_ and YPet-RIPK3_387-518_.

The different RHIM-containing components cannot be distinguished in these TEM images so we used confocal microscopy to visualise fibrils formed from mixtures of RHIM-containing fluorescent fusion proteins (Fig. 5b–e). We observed the formation of long fibrils that contained M45_1-90_ and either one or both of human RIPK1_497-583_ or RIPK3_387-518_. These assemblies are thioflavin T positive, confirming that these large heteromeric assemblies do have an amyloid cross-β substructure (Supplementary Fig. 4). Fibrils prepared from a mixture containing mCHERRY-M45_1-90_ and YPet-RIPK3_387-518_ were additionally imaged by super-resolution microscopy and transmission electron microscopy to provide higher resolution images. We found that the viral and host RHIMs were distributed throughout the fibrillar protein network formed by these proteins (Fig. 5f,g).

### The incorporation of M45 RHIM alters fibril stability

We reasoned that the incorporation of the viral M45 RHIM into host heteromeric fibrils might alter the properties of the fibrils, in addition to changing the morphology. Other studies of functional amyloid fibrils have noted that they are less resistant to treatment with the detergent sodium dodecyl sulphate (SDS) than some disease-associated amyloid fibrils^38,39^. We prepared homomeric and heteromeric assemblies from M45_1-90WT_ and M45_1-90AAAA_ with RIPK3_387-518_.. These assemblies were incubated with and without SDS, and then the preparations were analysed by agarose gel electrophoresis (AGE) (Fig. 6). This separated the material into (1) very large SDS-resistant fibrillar material that remained in the wells, (2) ThT-positive soluble amyloid oligomers and (3) monomeric protein. The M45_1-90WT_ and RIPK3_387-518_ mixtures contained soluble amyloid oligomers. The incorporation of the viral RHIM destabilised the complexes such that treatment with 2% SDS resulted in release of both constructs in monomeric form and, in the 1:1 mixtures, the generation of RIPK3-only fibrils. RIPK3-only fibrils were predominantly SDS-resistant and remained as large fibrils trapped in the wells (Fig. 6a). When the viral M45 RHIM was present in four-fold molar excess, little RIPK3-only fibrillar material remained in the wells. These results were supported by analysis of the soluble and insoluble material generated by dialysing mixtures of RHIM-containing constructs together, which showed that the presence of the M45_1-90WT_ fragment reduced the amount of RIPK3_387-518_ that was detected in the insoluble fraction (Supplementary Fig. 5). This activity of the M45 RHIM was dependent on the presence of the WT tetrad core sequence, as in the presence of M45_1-90AAAA,_ a large amount of SDS-resistant RIPK3-only material remained in the wells and very low levels of monomeric RIPK3_387-518_ were detected, even with four times as much M45_1-_ 90AAAA (Fig. 6a). The M45_1-277_ construct displayed a similar activity (Fig. 6b). It self-assembled into soluble homo-oligomers. It co-migrated with RIPK1_497-583_ but interacted strongly with RIPK3_387-518_ as judged by the narrow, intense orange-coloured co-localised band corresponding to soluble oligomers. In the absence of the intact RHIM core, M45_1-277AAAA_ formed large soluble oligomers with a wide mobility range but was unable to interact with RIPK1_497-583_ or RIPK3_387-518_, as judged by the presence of independently migrating oligomers of RIPK1_497-583_ and large RIPK3-only homo-oligomeric fibrils (Fig. 6b). The M45 RHIM is therefore able to compete with, and prevent, heteromeric RIPK1:RIPK3 RHIM-based amyloid formation and homo-oligomerisation of RIPK3 such as occurs after initial formation of the necrosome^40^.

**Figure 6:**
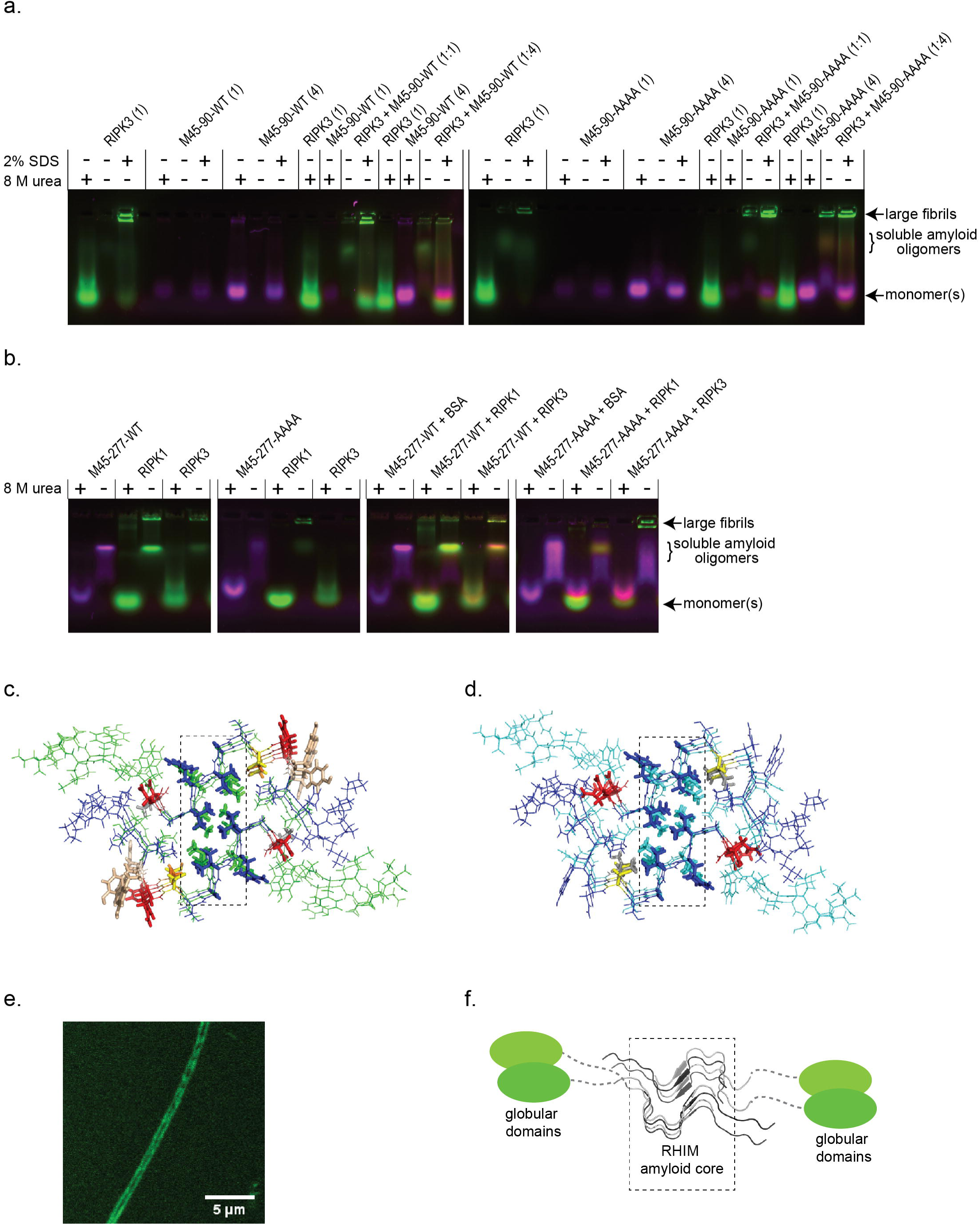
SDS AGE analysis of RHIM-based assemblies and model of the heteromeric M45:RIPK3 core. (**a**) SDS AGE analysis of YPet-RIPK3_387-518_ and mCHERRY-M45_1-90WT_ or mCHERRY-M45_1-90AAAA_ homomeric and heteromeric assemblies, formed by initial mixing in 8 M urea, followed by removal of urea by overnight dialysis. Dialysed samples were incubated with 0 or 2% SDS at RT for 10 min before electrophoresis. Monomeric forms of the protein constructs were maintained in 8 M urea prior to electrophoresis, as indicated. Protein components as indicated above each lane. (**b**) SDS AGE analysis of homomeric and heteromeric assemblies of YPet-RIPK1_497-583_, YPet-RIPK3_387-518_ and mCHERRY-M45_1-277WT_ or mCHERRY-M45_1-277AAAA_. BSA was included in M45-only samples to maintain the overall protein concentration constant. (**c**) Structure of the RIPK1:RIPK3 RHIM core. Figure prepared using PyMol Graphics System from PDB 5V7Z. The RIPK3 sequence is coloured in blue and RIPK1 is coloured in green. Tetrad residue side chains are shown as sticks; Tyr, Asn/Gln, Ser and Cys side chains are shown in wheat, red, grey and yellow, respectively. Core tetrad interactions across the opposing sheets are indicated by the dotted box. (**d**) Model of M45:RIPK3 RHIM core, prepared by appropriate mutation of RIPK1. The M45 sequence is coloured in cyan and RIPK3 in blue. Tetrad residue side chains are shown as sticks; Tyr, Asn/Gln, Val and Cys side chains are shown in wheat, red, grey and yellow, respectively. Core tetrad interactions across the opposing sheets are indicated by the dotted box. (**e**) Confocal microscopy image of a single fibril composed of YPet-M45. (**f**) Schematic representation showing associated globular domains (e.g. kinase or RNR, coloured green) flanking either side of the amyloid core formed by RHIMs (coloured grey).

## Discussion

These results demonstrate that the N-terminal 90 residues of M45 are sufficient to support interactions with the RHIMs of human RIPK1 and RIPK3 and to protect against TNFR-mediated necroptosis in human cells *in vitro*. The RHIM within the MCMV M45 protein is amyloidogenic. The ability of M45 to self-assemble into homomeric fibrils or to form heteromeric amyloid-structured complexes with RIPK1 and RIPK3 depends on the presence of the core tetrad sequence IQIG within the RHIM. This core sequence is also required for the ability of M45 to inhibit necroptosis^28^, strongly suggesting that the amyloid-based interactions made by M45 with RIPK1 and RIPK3 are associated with its anti-necroptosis activity. When probed with necroptosis-relevant RHIM-containing protein fragments, the viral RHIM preferentially interacts with the RIPK3 RHIM over the RIPK1 RHIM. In the presence of the M45 RHIM, a RHIM-encompassing region of RIPK3 is incorporated into metastable heteromeric fibrils rather than into SDS-resistant homo-oligomers.

The RIPK1:RIPK3 interaction and formation of the necrosome is central to TNFR-mediated responses if caspase-8 is inhibited by viral or chemical inhibitors^3^. We propose that the presence of M45 in cells results in the trapping of RIPK1 and RIPK3 within alternative heteromeric “decoy” amyloid structures and prevents the native RIPK1:RIPK3 RHIM-based amyloid interactions that are required to trigger cell death.

An important difference between RHIM-based amyloids and other disease-associated or most functional amyloids is the formation of heteromeric assemblies^30^. Heteromeric RHIM-based amyloid is central to necroptosis induction^11^. RNA-binding proteins containing low complexity sequences have the capacity to co-assemble into phase-separated droplets and hetero-fibrils, driven by intermolecular interactions between the prion-like domains, however the *in vivo* significance of heteromeric fibril assembly by these proteins is currently less clear than is the case for RHIM-based amyloid^38,39^. The multi-protein RHIM amyloid structures are similar to other higher-order assemblies such as signalosomes that form by polymerization and recruit multiple proteins with different activities^41^.

The key features of the heteromeric RIPK1:RIPK3 necrosome core have been revealed by ssNMR^32^. The RHIMs of RIPK1 and RIPK3 adopt a serpentine fold and form parallel β-sheets through intermolecular hydrogen binding. Two such sheets assemble to form a fibril through inter-tetrad and other interactions, which generate an oblong-shaped hydrophobic core (Fig. 6c). The structure is also stabilised by solvent-exposed side-chain-side-chain hydrogen bonds between stacked Asn and Gln residues, and Tyr and Ser/Thr side chains that form stacked ladder-like interactions on the periphery of the core.

We have used this structure to model the incorporation of the RHIM of M45. Since M45 and RIPK1 have the same core IQIG tetrad sequence, and we have determined that M45 preferentially forms heteromeric fibrils with RIPK3 over RIPK1, we have mutated the RIPK1 sequence to that of M45. These substitutions could be introduced into the complex structure without significant steric clashes (Fig. 6d). In the M45:RIPK3 RHIM model, the core tetrad interactions are unchanged from those in the RIPK1:RIPK3 structure but with the substitution with the viral protein, two Tyr and one Asn ladder are not possible. Instead, a new Asp-Asn side-chain interaction is likely and in the M45:RIPK3 structure, a Val-Cys ladder takes the place of the Ser-Cys ladder observed with RIPK1:RIPK3. Our model suggests that the M45:RIPK3 RHIM core will be similar to that formed by RIPK1:RIPK3 and hence it is likely that M45 mimics the host inter-RHIM interactions. The key role played by the tetrad residues in the hetero-fibrils is in line with our experimental observation that the AAAA mutation to M45 abolishes the ability of this protein to form a hetero-amyloid with the RHIM of RIPK3. Our experimental data demonstrate that the interactions driving M45:RIPK3 heteromeric assembly *in vitro* are stronger than those leading to M45:RIPK1 or RIPK1:RIPK3 assembly. Further mutagenesis studies could identify the M45-derived interactions responsible. The ssNMR-derived structure does not reveal heteromeric interactions between polypeptides that occupy the corresponding “rung” of opposing sheets. It is possible that the viral protein could be involved in such interactions with host partners. Interference by the viral protein with inter-sheet interactions would explain the branching and network morphology we observe in our heteromeric assemblies (Fig. 5a,f; Supplementary Fig. 2). The concentration of viral protein in the cell may also influence the make-up of the RHIM-based structures within cells, and hence control the final cellular outcome. The structure determined by Mompean *et al.* is consistent with confocal images we have collected of isolated M45_1-90WT_ homo-oligomeric fibrils which show fluorescent partner proteins along the two flanks of the fibril and a dark central core; an arrangement consistent with two opposed parallel β-sheets (Fig. 6e,f).

It is noteworthy that in our experiments the AAAA mutation of the core tetrad in RIPK3_387-518_ does not abolish self-assembly into an amyloid structure while this mutation in RIPK1_497-583_ and M45_1-90_ prevents self-assembly. This indicates that residues outside the tetrad of RIPK3 can support oligomerisation. These remain to be identified by further mutagenesis and structural studies of the entire RIPK3 RHIM. Although calculations based on the RIPK1:RIPK3 core structure suggest that RIPK3:RIPK3 homo-amyloid formation is less favourable than RIPK1:RIPK3 or RIPK1:RIPK1^32^, the formation of the RIPK1:RIPK3 necrosome is known to nucleate further polymerisation of RIPK3 that leads to activation of RIPK3 and interaction with MLKL^40^. The results of other studies show that the RHIM-based interactions of M45 with cellular partners prevent ZBP1-induced RIPK3 phosphorylation^7^ and autophosphorylation of RIPK3 when co-expressed in 293T cells^42^. HSV-1 and -2 encode RHIM-containing proteins, ICP-6 and ICP10, which have been shown to protect against necroptosis in human cells ^25,43^ and ICP10 expression is known to prevent phosphorylation of RIPK1 and RIPK3 compared to control^25^. The sequestration of host RHIM-containing proteins in heteromeric amyloids with viral proteins may therefore be general mechanism for modulation of host RHIM-based signalling by other RHIM-expressing viruses.

A different RHIM-based anti-necroptosis strategy is operated by some bacterial pathogens. Enteropathogenic *E. coli* (EPEC) is a gut pathogen that has been shown to encode a protease, EspL, that is delivered through the type III secretion effector system and which rapidly cleaves within the RHIM of RIPK1, RIPK3, ZBP-1/DAI and TRIF when these proteins are in a non -fibrillar form^44^. Amyloid fibril formation by the endogenous RHIM-containing proteins protects them from cleavage by EspL, allowing them to signal for necroptosis. Thus EspL appears to act by disabling the amyloid-based RHIM host defense pathways that would otherwise curtail infection^44^.

Several host RHIM:RHIM interactions occur in the absence of infection. An intact RHIM in RIPK1 suppresses undesirable ZBP-1:RIPK3 interactions during development that would lead to necroptosis^45,46^. The methods we have developed to probe and compare competing RHIM interactions can be applied to test the relative strengths of different RHIM:RHIM interactions and could shed light on the basis for cellular control of these potentially competing RHIM:RHIM interactions. Additionally, necroptosis is increasingly recognized in pathophysiological processes including ischaemia-reperfusion injury, and inflammation of the skin and intestinal epithelium^13,47^. Any development of therapies that target the necroptosis pathway will benefit from an understanding of the structural basis for viral inhibition of RHIM-based signaling.

## Acknowledgements

The work was supported by funds provided by Australian Research Council Discovery Project Grants DP150104227 and DP180101275 to M. Sunde. M. Steain was supported by National Health and Medical Research Council funding to Prof. Allison Abendroth and The University of Sydney Bridging Grant scheme. The authors acknowledge the facilities and the scientific and technical assistance of Sydney Microscopy & Microanalysis at the Australian Centre for Microscopy & Microanalysis at the University of Sydney. We thank Dr Louise Cole of the Bosch Institute Advanced Microscopy Facility for assistance with collection of confocal images, staff from the Bosch Institute Molecular Biology Facility for use of equipment, Charles Collyer for use of the X-ray generator, Sue McLennan for providing the HT-29 adenocarcinoma cells and James Murphy for providing the *E.coli* DNA gyrase subunit B. We thank Nicholas Della Marta, Monique Fischer, Laura Lau and Jake Campbell for preliminary experiments with RHIM peptides and cloning of fluorescent proteins. We thank Pascal Krotee and David Eisenberg for helpful discussions about the use of fluorescent proteins for studying RHIM interactions and models of heteromeric and homomeric amyloids, respectively.

## Supplementary Figures

**Supplementary Fig 1. The AAAA substitution of the core tetrad within the RHIM of M45_1-90_ abolishes its ability to self-assemble into fibrils.** Negatively stained EM of fibrils formed by (**a**) M45_1-90WT_-mCHERRY and **(b**) M45_1-90AAAA_-mCHERRY fibrils upon removal of 8 M urea by dialysis.

**Supplementary Fig 2. An increase in the amount of M45_1-90_ relative to RIPK3_387-518_ results in the formation of increasingly large fibrillar assemblies.** Negatively stained EMs of fibrillar assemblies formed from mixtures of M45_1-90WT_-mCHERRY and YPetRIPK3_387-518_ at (**a**) 1:1, (**b**) 2:1 and (**c**) 4:1 relative molar concentration. Monomeric forms of the proteins were mixed together and allowed to co-assemble upon removal of 8 M urea by dialysis.

**Supplementary Fig 3. The presence of an unrelated His_6_-ubiquitin fusion protein does not result in generation of fibrillar networks.** Samples of either (a) His-Ub-M45_1-90_ or (b) His-Ub-RIPK1_497-583_ RHIM fusion proteins were mixed with the non-RHIM-containing protein His-Ub-RodA in urea-containing buffer and allowed to assemble upon removal of urea by dialysis.

**Supplementary Fig 4. Heteromeric assemblies formed by RHIM-containing fragments are thioflavin T-positive, indicative of an amyloid structure.** mCHERRY-M45_1-90_ and YPet-RIPK3_387-518_ were allowed to coassemble during dialysis, following which thioflavin T was added and the dialysate imaged using a Cytation 3 Imager (BioTek). CFP, YFP and Texas Red filter sets were used to detect Thioflavin T fluorescence, YPet and mCHERRY respectively.

**Supplementary Fig 5. SDS-PAGE analysis of RHIM-containing assemblies resulting from overnight dialysis of individual constructs (a–c) or protein pairs (d–f).** RHIM-containing fusion proteins dialysed overnight to allow fibril assembly, either individually or in pairs after initial mixing in urea in the monomeric form. All RHIM constructs carried mCHERRY as fusion protein partner. RHIM-containing individual construct or pairs as indicated in each panel. The stability of the resulting material towards SDS treatment was tested. Samples are total dialysate (total), supernatant from centrifugation at 16000*g* for 10 min (soluble), pellet from centrifugation resuspended in 20 mM NaH_2_PO_4_, 150 mM NaCl, 0.5 mM DTT, pH 7.4 (insoluble), pellet from centrifugation treated with 2% SDS for 10 min at 37 °C and then separated again into supernatant (SDS-soluble) and pellet (SDS-insoluble). SDS-insoluble material was incubated in 8 M urea to solubilise material for detection by PAGE. This SDS analysis was performed once.

## Methods

### Expression of RHIM fusion proteins

RHIM-containing portions of mCMV M45 (Q06A28; residues 1-90), human RIPK1 (Q13546; residues 497-583), human RIPK3 (Q9Y572; residues 387-518), were produced as maltose binding protein (MBP) fusion proteins using the pMAL vector, as His_6_-Ubiquitin fusion proteins using the pHUE vector system (Catanzariti et al) or as His-tagged N- or C-terminal ECFP, YPet or mCHERRY fusion proteins using a vector prepared in-house. Successful cloning was confirmed by sequencing at the Australian Genome Research Facility at the Westmead Millennium Institute. Proteins were expressed in BL21(DE3) *E. coli* grown in LB media containing ampicillin at 37 °C for 2–3 hours, with induction by 0.5–1.0 mM IPTG when the OD_600 nm_ of the culture reached 0.6–was placed onto Parafil0m.8. All pMAL plasmids were grown in LB media containing 0.2% glucose to prevent amylase production. SDS PAGE analysis of cell lysates following induction of protein over-expression indicated that the MBP-RHIM fusion proteins were present in the soluble cell fraction while the His_6_-Ub-RHIM fusion proteins and His_6_-ECFP, His_6_-YPet or His_6_-mCHERRY fusion proteins were located in the insoluble fraction.

### Purification of MBP fusion proteins

Cells were lysed by incubation in 20 mM Tris.HCl, 200 mM NaCl, 1 mM EDTA and 0.5 mM DTT, pH 8.0 with lysozyme at 100 µg/mL, followed by a freeze thaw cycle. DNaseI and MgSO_4_ were added to final concentrations of 100 µg/mL and 100 mM respectively, and samples were incubated with stirring for 15 min at room temperature. Following the addition of AEBSF to 0.2 mM, samples were centrifuged at 4 °C at 10,000*g* for 40 min, to separate soluble material from insoluble cell debris. MBP fusion proteins were purified from the soluble fraction using amylose beads from New England Biolabs (MA, USA), as per the manufacturer’s instructions.

### Purification of His_6_-Ubiquitin fusion proteins

Cell pellets were lysed by suspension and incubation in 6 M GuHCl, 100 mM NaH_2_PO_4_, 20 mM Tris.HCl, 5 mM β-mercaptoethanol, pH 8.0, and soluble material further purified on Ni-NTA agarose (Life Technologies) under denaturing conditions, with exchange into 8 M urea, 100 mM NaH_2_PO_4_, 20 mM Tris.HCl, 5 mM β-mercaptoethanol at pH 6.0 for washing and pH 4.0 for elution from the Ni-NTA agarose.

### Purification of His_6_-ECFP, His_6_-YPet and His_6_-mCHERRY fusion proteins

Two methods for purification of polyhistidine-tagged fluorescent protein-RHIM fragment fusions were used during the course of this study. The first method was batch purification where cell pellets were lysed by suspension and incubation in 8 M urea, 100 mM NaH_2_PO_4_, 20 mM Tris.HCl, 5 mM β-mercaptoethanol, pH 8.0. Soluble material was purified using Ni-NTA agarose beads (Life Technologies) under denaturing conditions, with 8 M urea, 100 mM NaH_2_PO_4_, 20 mM Tris.HCl, 5 mM β-mercaptoethanol at pH 8.0 and in the presence of 20mM imidazole for washing steps and 300mM imidazole for elution from the Ni-NTA agarose. The second method involved using a His-trap (GE Healthcare) column. For this, cell pellets were lysed as described for the purification of MBP fusion proteins. The insoluble pellet was resuspended in 8 M urea, 100 mM NaH_2_PO_4_, 20 mM Tris, pH 8.0 and allow to dissolve by stirring for 2 h. The sample was centrifuged at 4 °C at 31,000*g* for 45 min, filtered and then purified on a 5mL His-Trap column (GE Healthcare) under denaturing conditions (8 M urea, 100 mM NaH_2_PO_4_, 20 mM Tris, pH 8.0 + 0.5mM DTT) using AKTA FPLC. The elution of protein was achieved with a gradient of imidazole. Protein elution was monitored by absorbance at 280 nm.

For both purification procedures, the eluted fractions were analysed by SDS-PAGE. Fractions containing the fusion proteins were pooled and concentrated using Amicon Ultra-15 Centrifugal Filter Units with MWCO 30kDa (Millipore). The concentration of the final concentrated protein sample was determined by BCA assay (Pierce).

### Removal of fusion tag (MBP and His_6_-Ubiquitin)

MBP-RHIM fusion proteins at concentrations of 1–4 mg/mL in 20 mM Tris.HCl, 200 mM NaCl, 1 mM EDTA, 2.5 mM CaCl_2_, 10 mM maltose pH 8.0 were incubated with 5 U Thrombin for 2 hours at 37 °C. Samples were centrifuged at 16 000*g* for 10 min to separate insoluble and soluble material and both fractions were analysed by SDS PAGE. His_6_-Ub-RHIM constructs in 8 M urea-containing buffer were dialysed overnight at room temperature, against 20 mM NaH_2_PO_4_, 50 mM NaCl pH 7.4 and then incubated with deubiquitylating enzyme UBP41 (produced in-house according to Catanzariti et al) at 37°C for 3 hours. The RHIM-containing fragments self-assembled to form insoluble material during dialysis and incubation with UBP41. Samples were centrifuged to separate the supernatant, containing the cleaved His_6_-Ubiquitin, from the pellet. The pellet containing the RHIM fragment was washed twice with 20 mM NaH_2_PO_4_, 50 mM NaCl pH 7.4, followed by addition of 6 M GuHCl, 100 mM NaH_2_PO_4_, 20 mM Tris.HCl pH 8.0 to solubilise the pellet and maintain the RHIM fragments in a monomeric form for storage.

### Size exclusion chromatography

MBP-RHIM fusion proteins, before and after treatment with enzyme to remove the fusion partner, were analysed by size exclusion chromatography using a Superdex 75 10/30 GL column (GE Healthcare Life Sciences), running in 20 mM NaH_2_PO_4_, 150 mM NaCl pH 7.4 with a flow rate of 0.4 mL/min. Protein elution was monitored by absorbance at 280 nm and peaks were analysed by SDS-PAGE and negative stain transmission electron microscopy.

### Congo red assays

Insulin (1 mg/mL) was incubated in 20 mM glycine pH 2.0, with shaking at 700 rpm for 4 hours at 60 °C to convert it into amyloid fibril form. Samples of monomeric insulin, insulin fibrils or MBP-RHIM fusion proteins, at a concentration of 100 µM, were prepared with 2 µM Congo red in 20 mM NaH_2_PO_4_, 150 mM NaCl pH 7.4 in a 1 mL cuvette. The absorbance of each protein-containing solution was measured over the wavelength range 400 to 650 nm and compared to the absorbance profile of a solution containing only Congo red in buffer.

### M45 RHIM peptide

A synthetic peptide encompassing the RHIM of M45 was purchased from Genscript with the sequence VRIMNGVSGIQIGNHNAMS. The peptide was solubilised in DMSO and then diluted with water to a final peptide concentration of 300 μM with addition of a trace amount of ammonium hydroxide to aid solubilisation. Addition of Thioflavin T indicated the development of a ThT-positive structure. Samples were analysed by transmission electron microscopy.

### Transmission Electron Microscopy (TEM)

A droplet of protein or peptide-containing solution (20 µL) was placed onto Parafilm™ and a carbon/formvar-coated copper grid (200 mesh, ProSciTech) was floated on the droplet surface for 1 minute. Excess solution was removed by touching the edge of the grid to filter paper and the grid was washed with filtered water three times and then stained by floating on a 20–µL droplet of 2% uranyl acetate for 2 minutes, followed by removal of excess solution by blotting. The grid was air-dried overnight at room temperature and then imaged using a Philips CM12 microscope operating at 120 kV. Digital images were recorded using a Morada 11 MegaPixel CCD camera camera and iTEM digital imaging system.

### ThT Kinetic Assays

His-Ub or fluorescent RHIM-containing fusion proteins in 8 M urea-containing buffer were diluted into phosphate buffer (25 mM NaH_2_PO_4_, 150 mM NaCl, 0.5 mM DTT, pH 7.4) containing 40 µM Thioflavin T to a final volume of 200 µL, protein concentration 10 µM His-Ub-M45, 2.5 µM mCHERRY-RIPK3_387-518_ or 2.5 µM YPet-RIPK1_497-583_ and residual urea concentration of 0.2 M. Three replicates of each protein sample were analysed simultaneously in a 96-well black fluorescence plate sealed with optically clear film (Corning^®^). Samples were incubated at 25 °C and fluorescence was measured at 480 nm, with excitation at 440 nm, every 60 s in a POLARstar Omega microplate reader (BMG Labtech).

### Fibril co-assembly for TEM

His_6_-Ub-RIPK3_387-518_ and RIPK1_497-583_ and M45_1-90_ were mixed together in 1:1 molar ratios or with an appropriate volume of urea-containing buffer and then dialysed against 25 mM NaH_2_PO_4_, 150 mM NaCl, 0.5 mM DTT pH 7.4 for a minimum of 4 hours. Samples were removed and used to prepare negatively stained grids for TEM, as above.

### Fluorescence anisotropy assays

Covalent labelling of RIPK1_497-583_ with the Alexa 488 fluorescence tag was performed in the amyloid state. Samples of 1–4 mL of 100 µM RHIM fragment in 6 M GuHCl, obtained from the cleavage of His_6_-Ub-RIPK1_497-583_, were dialysed against phosphate buffer at pH 8.0 with 0.5 mM DTT at room temperature for 3h, followed by further dialysis against 20 mM NaH_2_PO_4_, 150 mM NaCl pH 7.4 at pH 8.0 with two buffer changes. After the dialysis process, RIPK1-RHIM protein was in an amyloid state and the sample was centrifuged at 16 000g for 10 min. The supernatant was discarded and the amyloid pellet was resuspended in 1mL of 20 mM NaH_2_PO_4_, 150 mM NaCl, pH 8.0. A 50–µL volume of 10 mg/mL Alexa 488 (10 mg/mL in DMSO) was added to the amyloid sample. The sample was incubated on a rocker for 1 h at room temperature and then unbound dye was removed by centrifugation at 16000*g* for 10 min. The pellet was washed 3 times with 1 mL of 20 mM NaH_2_PO_4_, 150 mM NaCl, pH 8.0 and then resuspended in 1 mL of 8 M urea, 20 mM NaH_2_PO_4_, 150 mM NaCl, pH 7.4 for storage. The degree of labelling was calculated to be 5%. For polarization experiments, samples containing a molar ratio of 1:1999 of Alexa 488-RIPK1_497-583_ to unlabelled RIPK1_497-583_, RIPK3_387-518_ or M45_1-90_ in 20 mM NaH_2_PO_4_, 150 mM NaCl, pH 7.4 were prepared to a final volume of 300 µL. Total final protein concentrations of unlabelled protein were 0.85 µM for RIPK1, 0.69 µM RIPK3 and 1.05 µM for M45 and the residual urea concentration was 0.052 M. Three replicates of each protein sample were analysed simultaneously in a 96-well black fluorescence plate sealed with optically clear film (Corning^®^). Samples were incubated at 25 °C in a POLARstar Omega microplate reader (BMG Labtech) and excited at 480 nm. The parallel and perpendicular fluorescence emission was recorded at 520 nm, every 60 s. These fluorescence intensities were used to calculate the fluorescence polarization.

### X-ray fibre diffraction

His_6_-Ub-M45_1-90_ protein was allowed to assemble into fibrils by dialysis out of 8 M urea-containing buffer into water overnight. The insoluble fibrils were pelleted by centrifugation at 16000*g* for 10 min, resuspended in a small volume of water and allowed to air-dry from a droplet suspended between the wax-filled ends of two glass capillaries. X-ray fibre diffraction images were obtained using an in-house Cu Kα Rigaku rotating anode source (wavelength 1.5418 Å) and MARresearch image plate detector.

### SDS-stability assays

Protein samples (0.5–1 mL) of individual fluorescent RHIM-containing constructs as well as mixtures of these proteins were diluted to final individual protein concentrations of 20 µM with 8 M urea, 25 mM NaH_2_PO_4_, 150 mM NaCl, pH 7.4. Approximately 50 µL of each sample was removed for analysis as the monomeric, non-assembled control. The remaining solutions were dialysed overnight at room temperature against 25 mM NaH_2_PO_4_, 150 mM NaCl, 0.5 mM DTT, pH 7.4. Samples were removed and prepared with 4% glycerol and 0.0008% bromophenol blue and 0 or 2% SDS, incubated at room temperature for 10 min and then analysed by electrophoresis using a 1% agarose gel running in TAE buffer containing 0.1% SDS. Gels were imaged on a Bio-Rad Chemi-Doc imaging system with 605/50nm and 695/55nm emission filters. In order to study the stability of the amyloid fibrils by SDS-PAGE, mCHERRY RHIM-containing constructs, individually or in mixtures, were dialysed against phosphate buffer overnight, at room temperature. After dialysis, two 50–µL samples were centrifuged at 16,000*g* for 10 min and the resulting supernatants removed and pooled (soluble sample). One pellet was resuspended with 50 µL of 8 M urea, 25 mM NaH_2_PO_4_, 150 mM NaCl, pH 7.4 to allow visualisation of the insoluble material produced during dialysis (insoluble sample). The other pellet was resuspended in 50 µL of 2% SDS in 25 mM NaH_2_PO_4_, 150 mM NaCl, pH 7.4 and incubated at 37 °C for 10 minutes. SDS-containing samples were then centrifuged at 16,000*g* for 10 min. The supernatant was removed (SDS-soluble sample) and the pellet (SDS-insoluble sample) resuspended in 50 µL urea-containing buffer. All samples were boiled with NuPAGE LDS sample buffer and NuPAGE reducing agent (50 mM DTT, Thermo Scientific) and then analysed by SDS PAGE.

### Confocal microscopy

Protein samples (0.5 mL) in 8 M urea containing mixtures of monomeric, purified YPET, ECFP and mCHERRY RHIM-containing constructs at final individual protein concentrations of either 2.5 or 5 µM were dialysed overnight at room temperature against 25 mM NaH_2_PO_4_, 150 mM NaCl, 0.5 mM DTT, pH 7.4. Samples (15 µL) were placed on glass slides, protected with cover slips and imaged on a Zeiss LSM 510 Meta confocal microscope.

### Single molecule fluorescence spectroscopy

Mixtures of fluorescent fusion proteins with YPet or mCHERRY fluorophores were prepared at 70 µM in 8 M urea, 25 mM NaH_2_PO_4_, 150 mM NaCl, 0.5 mM DTT, pH 7.4 and then diluted 100– fold with 25 mM NaH_2_PO_4_, 150 mM NaCl, 0.5 mM DTT, pH 7.4 in a 192-well silicone plate for single molecule measurement before monitoring. Plates were analyzed at room temperature on a Zeiss Axio Observer microscope with a custom-built data acquisition setup. Fluorescence signal was collected and separated using a 565 nm dichroic mirror. For detection of YPET-tagged proteins, signal was passed through a 525/20 nm band pass filter; for detection of mCHERRY-tagged proteins a 580 nm long pass filter was used. For experiments with all three RHIM-containing proteins, protein mixtures containing YPet-RIPK3_387-518_, mCHERRY-RIPK1_497-583_ and His_6_-Ub-M45_1-90_ were prepared in 8 M urea-containing buffer, to provide samples containing the RIPK3:RIPK1:M45 RHIMs in the range ∼4:4:1 to ∼1:1:4. Samples were diluted 100-fold into 25 mM NaH_2_PO_4_, 150 mM NaCl, 0.5 mM DTT, pH 7.4 and 20 μL pipetted into a 192-well silicone plate for single molecule measurement. For experiments with multiple fluorophores, the signal from the two channels was recorded simultaneously in 1 ms time bins.

### Generation of M45-expressing HT-29 cells and cell death assays

M45_1-90_ and M45_1-277_ fragments were amplified from MCMV (Smith) and cloned upstream of the E.Coli DNA gyrase subunit B (residues 2-220) in the pCDH-MCS-EF1-neo lentivector. All insertions were validated by sequencing (Garvan Institute, Sydney, Australia). Lentivirus for M45_1-_ 90-gyrase, M45_1-277_-gyrase or empty vector control were made in 293T cells by co-transfecting the lentivectors with psPAX2 and pMD2G using Fugene HD (Promega). HT-29s were then transduced with the lentivirus particles in the presence of polybrene (Santa Cruz), and after 3 days G418 (Roche) was added for the subsequent 10 days to select for successfully transduced cells. M45_1-90_-gyrase, M45_1-277_-gyrase or empty-vector-control transduced HT-29s were seeded in 96 well plates (10^4^ cells/well) and allowed to adhere for 6 h before treating with 900 nM coumermycin A1 (Enzo) or media (no treatment control), followed 1 hour later with combinations of TNF (30 ng/ml, R and D), BV-6 (1 μM, Selleckchem) and z-VAD-fmk (25 μM, R and D), or DMSO only (Sigma-Aldrich) as control. Cell viability was then assessed 17-18 hour post-treatment by measuring levels of intracellular ATP, using the CellTitre-Glo2 Assay (Promega). Data was expressed as percentage of cell survival relative to the DMSO only control. Luminescence was measured using an Infinite M1000 Pro plate reader (TECAN).

